# Age-dependent impairment of disease tolerance is associated with a robust transcriptional response following RNA virus infection in *Drosophila*

**DOI:** 10.1101/2020.09.21.307017

**Authors:** Lakbira Sheffield, Noah Sciambra, Alysa Evans, Eli Hagedorn, Megan Delfeld, Casey Goltz, Janna L. Fierst, Stanislava Chtarbanova

**Affiliations:** Department of Biological Sciences, University of Alabama, 300, Hackberry lane, Tuscaloosa, AL-35487, USA

**Keywords:** aging, infection, tolerance, RNA virus, *Drosophila melanogaster*, transcriptomics

## Abstract

Advanced age in humans is associated with greater susceptibility to and higher mortality rates from infections, including infections with some RNA viruses. The underlying innate immune mechanisms, which represent the first line of defense against pathogens, remain incompletely understood. *Drosophila melanogaster* is able to mount potent and evolutionarily conserved innate immune defenses against a variety of microorganisms including viruses and serves as an excellent model organism for studying host-pathogen interactions. With its relatively short lifespan, *Drosophila* also is an organism of choice for aging studies. Despite numerous advantages that this model offers, *Drosophila* has not been used to its potential to investigate the response of the aged host to viral infection. Here we show that in comparison to younger flies, aged *Drosophila* succumb more rapidly to infection with the RNA-containing Flock House Virus (FHV) due to an age-dependent defect in disease tolerance. In comparison to younger individuals, we find that older *Drosophila* mount larger transcriptional responses characterized by differential regulation of more genes and genes regulated to a greater extent. Our results indicate that loss of disease tolerance to FHV with age possibly results from a stronger regulation of genes involved in apoptosis, activation of the *Drosophila* Immune deficiency (IMD) NF-kB pathway or from downregulation of genes whose products function in mitochondria and mitochondrial respiration. Our work shows that *Drosophila* can serve as a model to investigate host-virus interactions during aging and sets the stage for future analysis of the age-dependent mechanisms that govern survival and control of virus infections at older age.

## Introduction

Infectious diseases, including viral infections, represent an important burden among the elderly. For instance, older age is a major risk factor for increased morbidity and mortality to numerous viral pathogens including the Severe acute respiratory syndrome (SARS) associated coronavirus-2 (SARS-CoV-2), the agent responsible for the current COVID-19 pandemic (Nikolich-Zugich *et al*. 2020). Immunosenescence, a collective term used to describe the progressive functional decline of the immune system over time, is associated with the increased susceptibility to infections and lower responsiveness to vaccination observed in the elderly (Leng and Goldstein 2010). Considerable progress has been made in understanding how aging affects both, the innate and adaptive immune systems, however, the causes underlying immunosenescence remain incompletely elucidated. In particular, the age-dependent mechanisms leading to dysregulated innate immunity, which represents the first line of defense against invading pathogens, are less well documented (reviewed in (Nikolich-Zugich 2018)). Moreover, the exact factors and molecular events contributing to the more rapid death of the aged organism following virus infection are not fully understood (Nikolich-Zugich *et al*. 2020). With an increasing aging population (He *et al*. 2016), there is a great need to further our knowledge of the mechanisms underlying the capability of the aged organism to survive infection to ensure appropriate preventive and treatment strategies and to improve healthspan.

Pioneering research using the genetically tractable model organism *Drosophila melanogaster*, which in contrast to vertebrates is devoid of a classic adaptive immune system, has uncovered conserved mechanisms of activation of innate immunity in response to bacterial and fungal pathogens. Following bacterial or fungal infection, two nuclear factor kappa B (NF-κB) pathways, Toll and immune deficiency (IMD), which share similarities with mammalian Toll-like receptor/interleukin (IL)-1 receptor and tumor necrosis factor receptor (TNFR) pathways, respectively, are activated and mediate the transcription of downstream targets including antimicrobial peptides (AMPs) (reviewed in (Lemaitre and Hoffmann 2007)). *Drosophila* also detect and respond to viral pathogens via multiple mechanisms that mediate antiviral defenses. RNA interference (RNAi), which relies on production of virus-derived small interfering RNAs (siRNAs), provides broad protection against RNA and DNA viruses. Cellular processes such as apoptosis, apoptotic bodies’ clearance by plasmatocytes (macrophage-like cells in *Drosophila*) and autophagy also represent effective antiviral mechanisms (reviewed in (Mussabekova *et al*. 2017) and in (Lamiable and Imler 2014)). In *Drosophila*, viral infections are also associated with complex transcriptional responses that reflect the regulation of cellular pathways, production of cytokines and effector molecules, changes in stress response and physiology (reviewed in (Mussabekova *et al*. 2017)). Although the *Drosophila* genome does not encode for interferon genes, the protein encoded by the stimulator of interferon genes (STING), which in mammals activates NF-κB and interferon signaling in response to viral infection, is present in this organism. dSTING recently was shown to contribute to antiviral immunity by interacting with some of the components of the *Drosophila* IMD pathway in response to picorna-like viruses (Goto *et al*. 2018) and by activating downstream autophagy in response to ZIKA virus infection in the brain (Liu *et al*. 2018). In addition to these mechanisms that are in control of pathogen burden (also referred as resistance mechanisms), the outcome of infection is determined by the ability of the host or to endure the damaging effects caused by the pathogen or resulting from immunopathology (a phenomenon known as disease tolerance). Both resistance and tolerance are considered components of host immunity and effective tolerance mechanisms allow resistance mechanisms to operate in a more optimal way (Martins *et al*. 2019).

Aging in *Drosophila* also leads to deregulation of innate immunity. For instance, expression of several genes encoding AMPs downstream of NF-κB pathways increases with age (reviewed in (Garschall and Flatt 2018)), similar to inflammaging, the low grade chronic inflammation that accompanies aging (Franceschi *et al*. 2000) in mammals. Additionally, the phagocytic capacity of *Drosophila* macrophages declines with age (Mackenzie *et al*. 2011; Horn *et al*. 2014). Aged *Drosophila* also are more sensitive to infections with Gram-negative bacteria, Gram-positive bacteria, fungi and viruses such as *Drosophila* C virus (DCV) and the Flock House virus (FHV) (Ramsden *et al*. 2008; Eleftherianos *et al*. 2011; Fabian *et al*. 2018). However, there is still a very limited understanding of how antiviral immunity operates as a function of age in *Drosophila*. With increasing evidence for impaired defenses against viruses in the aged organism, flies can serve as a prime genetic model of aged host-virus interactions and can offer unique opportunities for mechanistic dissection of age-dependent innate immune responses.

In the present study, we conducted comparative analysis of survival, virus load and gene expression between young and aged *Drosophila* following infection with the Flock House Virus (FHV). FHV is a small, non-enveloped virus, whose genome is composed of two positive, single-stranded RNA molecules (Venter and Schneemann 2008). We report that older flies succumb faster to FHV infection without accumulating higher virus loads, suggesting that a tolerance mechanism becomes impaired with age. Additionally, we show that aged flies mount a more robust transcriptional response to FHV than young flies, including the regulation of innate immunity genes; response, which is different from the response of flies undergoing aging. Genes encoding components of the apoptotic process are predominantly regulated in aged, FHV-infected flies. Additionally, we show that several genes whose gene products function in mitochondria and mitochondrial respiratory chain are specifically downregulated in aged, FHV-infected flies. We also demonstrate that among genes that do not belong to specific gene ontology categories, the expression of several encoding for non-coding RNAs (ncRNAs) changes in aged, FHV-infected flies in comparison to young, FHV-infected *Drosophila* and flies undergoing aging. Collectively, our work shows that virus infection in aged flies triggers profound changes in transcriptomics and establishes *Drosophila* as a model that allows investigation of the age-dependent mechanisms underlying the response and survival to viral infection.

## Results

### FHV infection leads to decreased survival but not increased virus load in aged

#### Drosophila

To determine how age affects survival to infection with FHV, we injected 5-day old and 30-day old wild type (*OregonR*) male or female flies with either Tris buffer (control) or FHV and recorded their survival every 24 hours. 30-day old flies (median survival = 6.75±0.17 days for males and 7.17±0.28 days for females) showed decreased survival in comparison to 5-day old flies (median survival = 8.17±0.15 days for males and 8.04±0.23 days for females) (Figure 1A and Figure S1A). We hypothesized that the higher mortality in aged flies could result from increase in FHV load. To test this, we separately injected groups of male and female 5- and 30-day old flies with either Tris or FHV and measured virus load 96h post infection (p.i.) using quantitative reverse transcription PCR (RT-qPCR). For both sexes we observed comparable, non-significantly different levels of FHV RNA1 (*FHV1*) expression between young and aged flies (Figure 1B). Interestingly, although survival curves overlapped at 5 days of age between both sexes (Figure 1A and Figure S1A), virus load was significantly lower in females in comparison to males (Figure 1B). At 30 days of age, females showed significant, two-fold decrease in virus load in comparison to males (Figure 1B), which was accompanied with slightly better, although non-significantly different median survival to FHV (6.75±0.17 days for males and 7.17±0.28 days for females, Figure S1A). In support of the data obtained for *OregonR* male flies, similar differences in survival between 5- and 30-day old flies, and comparable *FHV1* load at 72h p.i. between animals of the two age groups was observed for males of another genotype, *y*_*1*_ *w*_*67c23*_ (Figure S1B). Additionally, we found non-significant differences between FHV titers in circulating hemolymph (insect blood) of 5- and 30-day old female *w*_*1118*_ flies 96h p.i. (Figure S1C).

**Figure 1.**
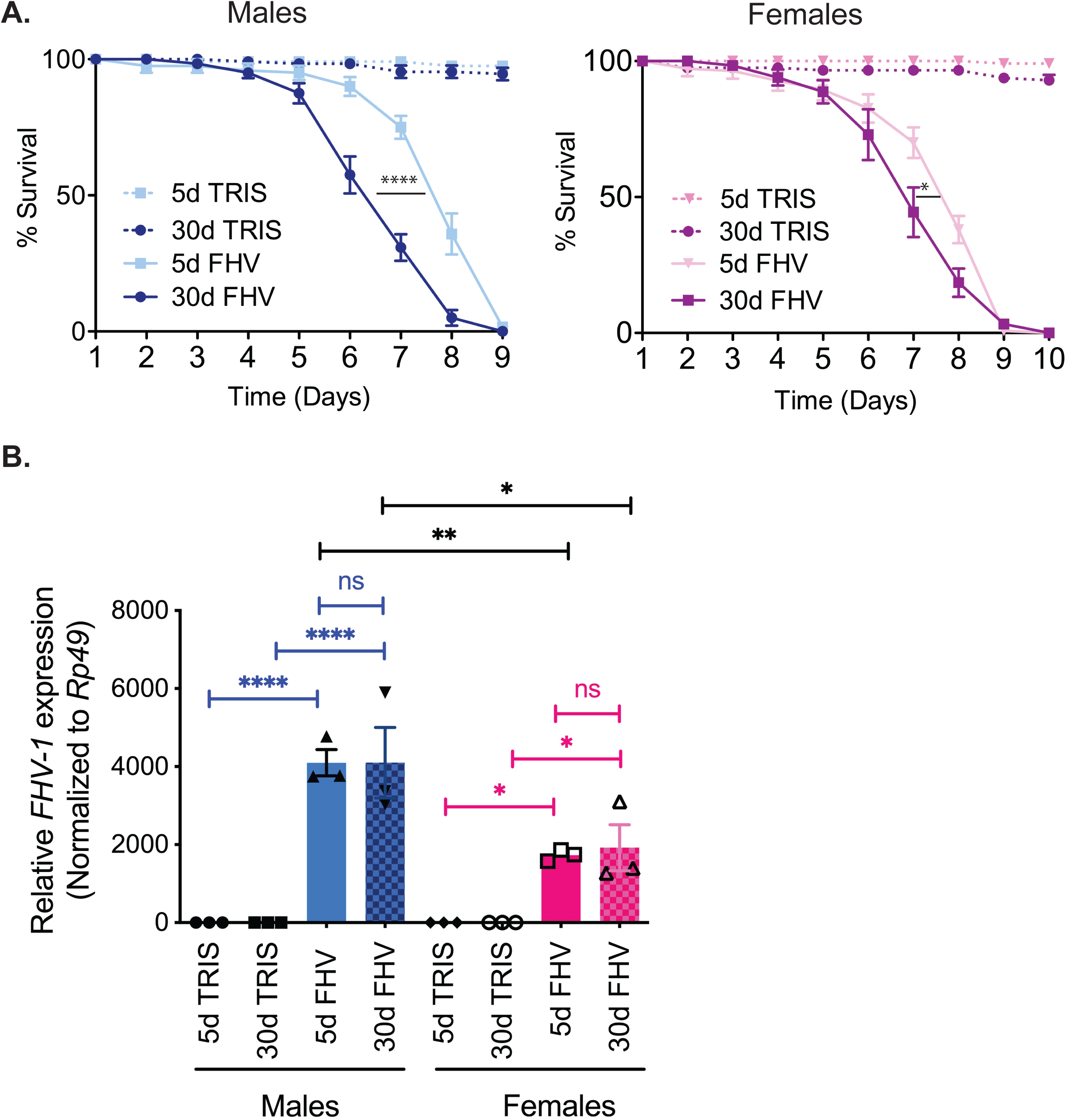
FHV infection triggers more rapid death of aged *Drosophila* without accumulation of higher virus load. **A**. Survival curves of young and aged male and female *OregonR Drosophila* that have been infected with FHV or control-injected with the same volume of Tris. Statistics of median survival to FHV are based on a Student’s *t*-test. ****: *p*<0.0001, *: *p*<0.05. **B**. Virus load determined by *FHV RNA1* expression reveals comparable titers between young and aged animals. Note that significant difference is observed between males and females in both young and aged flies. Statistics are based on two-way ANOVA followed by Tukey post-test to correct for multiple comparisons. ****: *p*<0.0001, ** : *p*<0.001, *: *p*<0.05, ns=non-significant.

Altogether, these results indicate that 30-day old *OregonR* flies succumb faster to infection with FHV than younger flies, where males exhibit higher mortality in comparison to females. Moreover, decreased survival of infected 30-day old flies is not accompanied with increased FHV titers in whole flies, suggesting that the aged organism is able to control viral pathogen burden.

### Aged *Drosophila* mount a robust transcriptional response following FHV infection

Both aging and virus infection lead to changes in the *Drosophila* transcriptome (Pletcher *et al*. 2002; Kemp *et al*. 2013; Chtarbanova *et al*. 2014). We hypothesized that aged flies infected with FHV mount a distinct transcriptional response in comparison to young flies, potentially accounting for the observed increase in mortality. To test this, we performed transcriptomics analysis using RNA sequencing (RNA-Seq) on 7-day old (young) and 25-day old (aged) male *OregonR Drosophila* injected with either Tris or FHV at 24h and 48h following injection. This sex was chosen because aged males showed more pronounced effect on survival than females. The time points were chosen early in the infection process before differences in survival between age groups were detected. As an additional control, we used non-infected young and aged flies to control for the effects of aging alone in absence of infection. An average of 95.4% of each RNA-Seq library (Table S1) aligned to the *D. melanogaster* genome (Table S2). We validated the RNA-Seq data for aging and the 48h post FHV infection time point using specific primers and RT-qPCR analysis for four genes per experimental condition. We confirmed that in aging flies *Cpr67Fb* and *CG15199* were upregulated and *Acp54A1* and *Lman III* were downregulated. In young *Drosophila*, 48h after FHV infection, *Upd2* and *Ets21c* were upregulated and *Rfabg* and *Diedel 3* were downregulated in comparison to Tris-injected controls. In aged flies, FHV infection led to upregulation of *Or85a* and *Upd3* and downregulation of *IM14* and *GNBP-Like 3* (Figure S2).

To evaluate the overall similarity and differences between treatments, we used principal component analysis (PCA). PCA creates new orthogonal variables that maximize the variance observed in high dimensional space, in this case the expression levels across annotated *D. melanogaster* genes. Principal component 1 (PC1) explains the majority of variance across samples and treatments while subsequent PCs explaining the next most uncorrelated variance. We observed that both young and aged FHV samples displayed a composite signature of gene regulation fundamentally different from non-infected young, non-infected aged or Tris-injected samples (Figure 2A). PC1 explained 54% of the variance in gene expression and separated young and older FHV-infected hosts from non-infected controls. PC2 explained 20% of the variance in expression and further separated FHV-infected hosts 24h and 48h p.i. Each of the three replicates grouped by treatment with the exception of the young Tris-injected flies 24h and 48h post-injection, which overlapped.

**Figure 2.**
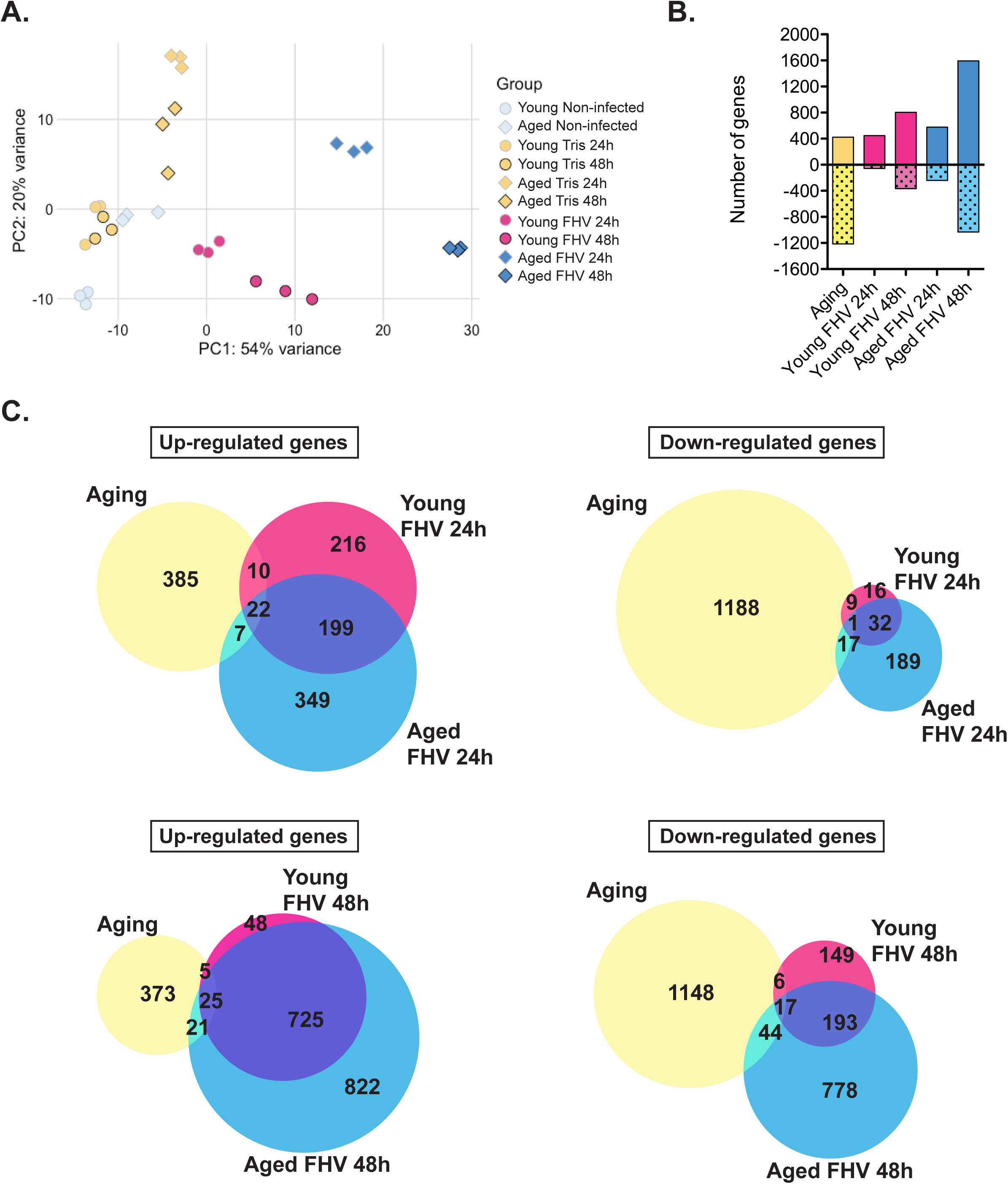
FHV infection of aged flies leads to a robust transcriptional response. **A**. Principal component analysis for all experimental samples. **B**. Comparison of the number of differentially regulated genes at least two-fold in all conditions. Positive values represent upregulated genes and negative values represent downregulated genes. **C**. Venn diagrams showing overlaps between differentially regulated genes for selected experimental conditions.

Differential gene expression analysis following FHV infection revealed that more genes were significantly regulated (*p adj* < 0.05) at least two-fold at 48h p.i. in comparison to 24h p.i. in both age groups. More genes were differentially changed in aged FHV-infected flies in comparison to young flies for both time points (Figure 2B, Table S3). Overall, in young flies, the expression of 505 genes was differentially regulated 24h p.i. vs 1,168 genes 48h p.i. In aged flies, we observed differential regulation of 816 genes at 24h p.i. and 2,625 genes at 48h p.i. The process of aging itself differentially regulated expression of 1,639 genes (Figure 2B). We note that in aging flies, more genes are downregulated than upregulated, whereas in aged, FHV-infected flies there are fewer downregulated than upregulated genes (Figure 2B).

Among the genes differentially regulated during aging, we observed a very small overlap with genes regulated by infection at either young or older age, 24h or 48h p.i. (1.4% and 2.5%, respectively) (Figure 2C). At 24h p.i., ∼50% of upregulated genes and 57% of downregulated genes in young flies overlapped with genes upregulated in aged, FHV-infected flies. At 48h p.i. in young flies, 93% of upregulated genes overlapped with upregulated genes in FHV-infected aged flies and 57% of downregulated genes overlapped between the two age groups (Figure 2C).

Altogether, these results indicate that aged male flies mount a larger transcriptional response following FHV infection than younger flies, a signature that is different from the transcriptional changes taking place during the aging process itself. The fact that most of commonly regulated genes between young and aged FHV-infected flies were found to overlap as a function of time (86% of up- and 87% of down-regulated genes, Figure S3), is in support of the hypothesis that the age-dependent defect in disease tolerance is unlikely to result from the regulation of these genes. Rather, our data suggest that impaired tolerance in aged flies could be due to differential regulation of the genes that are uniquely expressed in infected young flies, uniquely expressed in infected aged flies or a combination of both.

### FHV infection triggers transcriptional changes in similar and different biological processes in young and aged *Drosophila*

To visualize biological processes regulated by aging and FHV infection in young and aged flies, we performed gene ontology (GO) analysis. The number of genes with Flybase ID (FBgn number) without a matching DAVID ID is listed in Table S4. We note that most differentially regulated genes with a DAVID ID were labeled as “Others” (Figure S4). For instance, 76% of differentially regulated genes for the Aging group did not match a specific biological process. For Young FHV24h, Young FHV48h, Aged FHV24h and Aged FHV48h, these percentages are 53%, 59%, 60% and 59%, respectively (Figure S4).

Our GO analysis revealed a complex signature. For instance, aging led to changes in expression of genes belonging to 57 biological processes. Five of them (‘defense response’, ‘response to bacterium’, ‘antibacterial humoral response’, ‘defense response to Gram-positive bacterium’ and ‘oxidation-reduction’) overlapped between all five experimental conditions. ‘Mannose metabolic process’ and ‘protein refolding’ were in common between Aging and Aged FHV24h groups and ‘sperm storage’ between Aging and Aged FHV48h groups. Processes identified in common between the Aging group and young and aged FHV-infected flies were ‘circadian rhythm’, ‘multicellular organism reproduction’ and ‘proteolysis’ (Figure 3 and Table S5). In *Drosophila*, aging leads to both, deregulation of organismal reproduction (Tatar 2010) and innate immunity (Pletcher *et al*. 2002; Zerofsky *et al*. 2005; Kounatidis *et al*. 2017). In flies, it is also well established that physiological trade-offs exist between immune activation and reproductive capacity (Zerofsky *et al*. 2005), potentially accounting for the differential regulation of genes involved in organismal reproduction after FHV infection in both, young and aged flies. Among the 46 biological processes specific to Aging, we find genes belonging to ‘metabolic process’ and ‘spermatogenesis’ GO categories (Figure 3). This aligns with previous studies showing that aging impacts male germline stem cells and leads to decrease in spermatogenesis (Boyle *et al*. 2007) as well as with previous observations that the aging process leads to differential regulation of genes involved in *Drosophila* metabolism (Pletcher *et al*. 2002).

**Figure 3.**
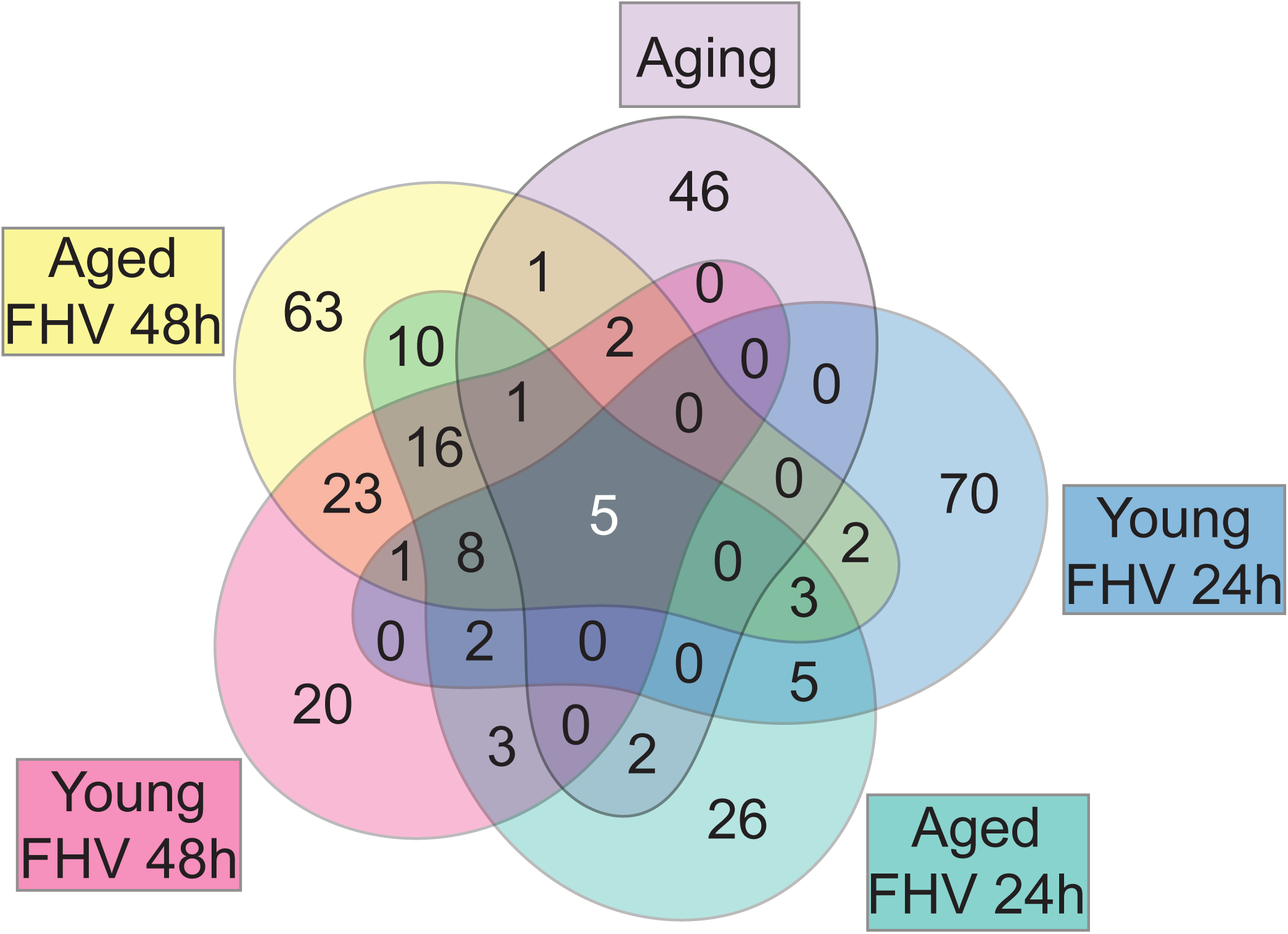
Common and distinct biological processes are regulated by aging and FHV infection in *Drosophila*. Venn diagram showing overlaps between Biological processes among different experimental groups, based on Gene ontology analysis.

At 24h p.i., we identified more biological processes in young flies than in aged animals (96 vs 80, respectively), among which five overlapped between the two age groups. 70 and 26 biological processes were specific to Young FHV24h and Aged FHV24h, respectively (Figure 3 and Table S5). At 48h p.i., we found an opposite trend with 81 and 135 biological processes in young and aged flies, respectively, among which 23 overlapped. We found 20 and 63 biological processes to be specific to the Young FHV48h and Aged FHV48h groups, respectively (Figure 3 and Table S5).

In both young and aged flies, FHV infection led to differential regulation of genes involved in processes associated with the nervous system. Clustering analysis identified one module of ‘neurogenesis’ genes that were strongly upregulated in the Aged FHV48h group and regulated to a lesser extent in Young FHV48h and Aged FHV24h groups (Figure S5A). For instance, among genes belonging to this GO category at 48h p.i., the gene *midlife crisis* (*mdlc*), which is required for neuroblast proliferation and neuronal differentiation in *Drosophila* (Carney *et al*. 2013), was upregulated to a greater extent in aged flies. *Ankyrin repeat and LEM domain containing 2* (*Ankle2*), the *Drosophila* ortholog of human ANKLE2, which is a target of the ZIKA virus NS4 protein (Shah *et al*. 2018), also showed stronger upregulation in aged flies (Figure S5B and Table S3). Other biological processes linked to the nervous system development and function for which genes were enriched in young and aged FHV-infected groups were ‘lateral inhibition’, ‘sleep’ and ‘ventral cord development’ (Table S5). The significance of this regulation is not known as FHV has not been previously demonstrated to target the nervous system, but rather the *Drosophila* heart and fat body (Eleftherianos *et al*. 2011). Among other common processes identified we find ‘innate immune response, ‘protein folding’, ‘rRNA processing’ and ‘response to heat’. In *Drosophila*, the heat shock response plays an antiviral role against the RNA viruses DCV and Cricket Paralysis Virus (CrPV), as well as against the DNA Invertebrate Iridescent Virus 6 (IIV-6) (Merkling *et al*. 2015). Indeed, several heat shock proteins belonging to the biological process ‘response to heat’ were upregulated in both young and aged FHV-infected flies (Table S3 and Table S5). This suggests that following FHV infection, this branch of antiviral immunity is functional in aged flies.

Interestingly, genes belonging to additional categories associated with nervous system’s function such as ‘neuromuscular synaptic transmission’, ‘transmembrane transport’ and ‘neurotransmitter secretion’ were specifically found in the Young FHV24h group. On the other hand, among processes specific to Aged FHV24h we found ‘autophagic cell death’, and ‘regulation of autophagy’ (Table S5). Among processes specifically enriched 48h p.i., we found ‘regulation of transcription, DNA-templated’, ‘transmembrane receptor protein tyrosine kinase signaling pathway’ and “protein ubiquitination” in young flies and ‘phagocytosis’, ‘programmed cell death’ and ‘peptidoglycan recognition protein signaling pathway’ in aged flies. The latter category contained multiple genes encoding for components of the *Drosophila* IMD pathway. Finally, among the processes specifically regulated in aged flies at both 24h and 48h p.i., we found ‘apoptotic process’, ‘determination of adult lifespan’ and ‘chromatin remodeling’ (Table S5).

Overall, these results indicate that despite a large number of “other” genes, genes belonging to identifiable common and distinct categories of biological processes are regulated by aging and FHV infection of young and aged flies. Although our results identify specific categories of biological processes for each experimental group (Figure 3), at this stage we are not able to determine whether the age-associated impairment of tolerance depends on the regulation of genes that are specifically regulated in young or/and aged flies.

### Profiles of innate immunity gene expression are distinct between aging and FHV infection in young and aged flies

We found genes belonging to ‘innate immune response’ to be differentially regulated in both young and aged FHV-infected flies at 24h and 48h p.i. Clustering analysis for this GO category identified similar patterns of differential gene expression between young and aged, FHV-infected flies. However, this regulation was to a greater extent in aged, FHV-infected flies (Figure 4A). The expression pattern of immunity-related genes during aging, for most part, was opposite than following FHV infection. Consistent with previous reports, we observed increased expression of several AMP and Immune induced molecule (IM) genes (*CecA1, Def, IM3, Drs, IM2, IM1, IM4, IM14* and *IM33*) as well as *GNBP-like 3* in aging flies (Figure 4A, Figure S6 and Table S3). Interestingly, in both young and aged flies, FHV infection led to strong downregulation of most AMP and IM genes, despite a robust upregulation of the mRNA encoding the NF-κB factor Relish (Figure 4A and Table S3). In aged FHV-infected flies, we observed marked upregulation of IMD pathway components *PGRP-LE, imd, key* (*IKKγ*) and *AttD*. This upregulation was to a greater extent in the Aged FHV48h group (Figure 4A and Figure S6). In comparison to aging and young FHV-infected *Drosophila*, we found *dSTING*, whose product acts upstream of Relish to protect flies against infection with DCV and CrPV (Goto *et al*. 2018), to be strongly upregulated in aged FHV-infected flies (Figure S6). In comparison with non-infected flies, Tris injection alone affected to a greater extent the expression in aged flies of several *Turandot* (*Tot*), IM and AMP genes at the 24h time point (Figure 4A). This suggests that older animals respond to injury by upregulating innate immunity genes toa greater extent than younger flies.

**Figure 4.**
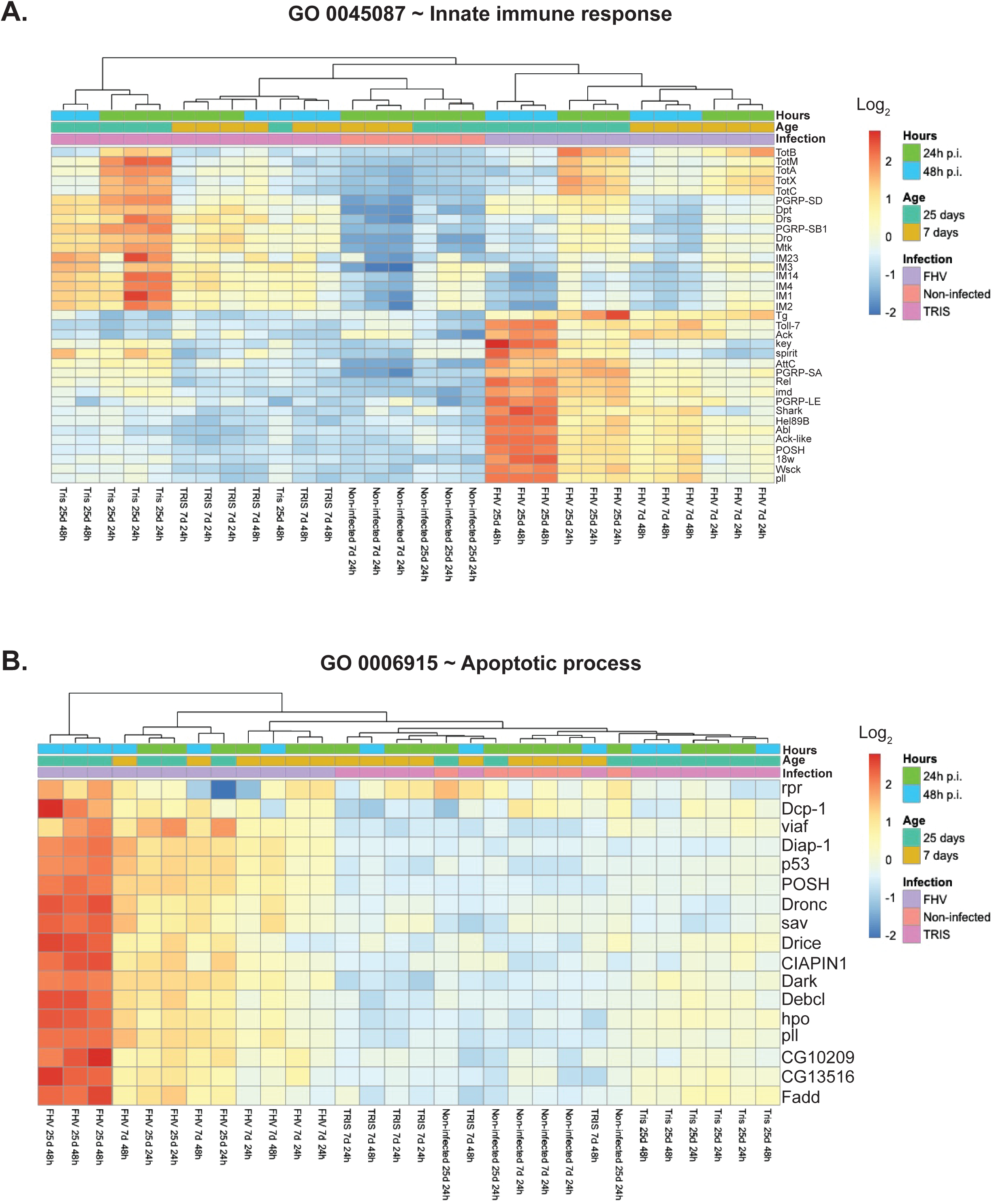
Regulation of innate immunity and programmed cell death genes by aging and FHV infection. **A**. Heatmap comparing the expression of genes belonging to the GO category ‘innate immune response’ based on their expression in all experimental groups. **B**. Heatmap comparing the expression of genes belonging to the GO category ‘apoptotic process’ based on their expression in all experimental groups.

Altogether, these results indicate that aging and virus infection lead, for the most part, to different transcriptional signatures for innate immunity genes. Additionally, aged flies carry out an overall stronger response to FHV than younger flies and regulate expression of more components of the IMD pathway. Because overactivation of the IMD pathway exerts detrimental effects on *Drosophila* tissues and leads to premature death (Cao *et al*. 2013; Kounatidis *et al*. 2017), our results also suggest that the specific upregulation of components of this pathway could be responsible for impaired tolerance and decreased survival in aged FHV-infected flies.

### Strong apoptotic gene expression signature in aged *Drosophila* following FHV infection

“Apoptotic process” was among the GO categories represented specifically in aged FHV-infected flies (Figure 3 and Table S5). Apoptosis is a form of programmed cell death and p53-dependent early induction of pro-apoptotic genes has been proposed to play a protective role against FHV infection in *Drosophila* (Liu *et al*. 2013). Additionally, infection with FHV of *Drosophila* cells in culture leads to induction of apoptosis, which is dependent on the effector caspase DrICE, the initiator caspase Dronc and its cofactor Dark (Settles and Friesen 2008). Consistent with this, we find that FHV infection in young and predominantly in aged flies leads to transcriptional changes in expression of several genes involved in the apoptotic process (*p53, Dronc* and *Dark*) 48h after FHV infection (Figure 4B). In cell culture, over the course of FHV infection, protein levels of the *Drosophila* inhibitor of apoptosis (Diap-1) are progressively depleted as a possible result of host cell translational shut down (Settles and Friesen 2008). In our RNA-Seq data we find that *diap-1* mRNA increased post infection and to higher levels in aged flies in comparison to young adults (Figure 4B). This change could potentially represent a compensatory increase in *diap-1* mRNA as a result of the rapid depletion of the protein. Aged, FHV-infected flies also exhibited increased expression of additional genes belonging to this GO category (Figure 4B).

Collectively, these results are in agreement with previous findings that FHV infection leads to apoptotic cell death and suggest that either more rapid or widespread activation of cell death takes place in the aged, FHV infected organism.

### Genes specifically downregulated in aged flies following FHV infection are enriched for metabolism and mitochondria

We performed pathway enrichment analysis on the genes specifically regulated in the Aged FHV48h group (822 upregulated and 778 downregulated genes, respectively; Figure 2C) using the Kyoto Encyclopedia of Genes and Genomes (KEGG) database. We identified ‘purine metabolism’, ‘pyrimidine metabolism’ and ‘RNA polymerase’ for upregulated genes, likely reflecting the higher transcriptional rates observed in aged animals in comparison with younger adults following FHV infection. Among downregulated genes we found ‘metabolism’ as a strongly enriched category (Figure S7). Using cellular component GO analysis, we found among downregulated genes several belonging to mitochondria and to mitochondrial electron transport chain (ETC) complexes I, III and IV (Figure 5). Mitochondrial ETC complexes represent a series of four protein complexes (I-IV) distributed along the inner mitochondrial membrane, where they function to pump protons from the mitochondrial matrix into the intermembrane space. ETC complexes are coupled to Complex V (the ATP synthase), which helps the production of ATP (Burke 2017). The observed downregulation of these genes could reflect a virus induced defect of the mitochondrial respiratory chain affecting ATP levels, specifically in the infected aged organism.

**Figure 5.**
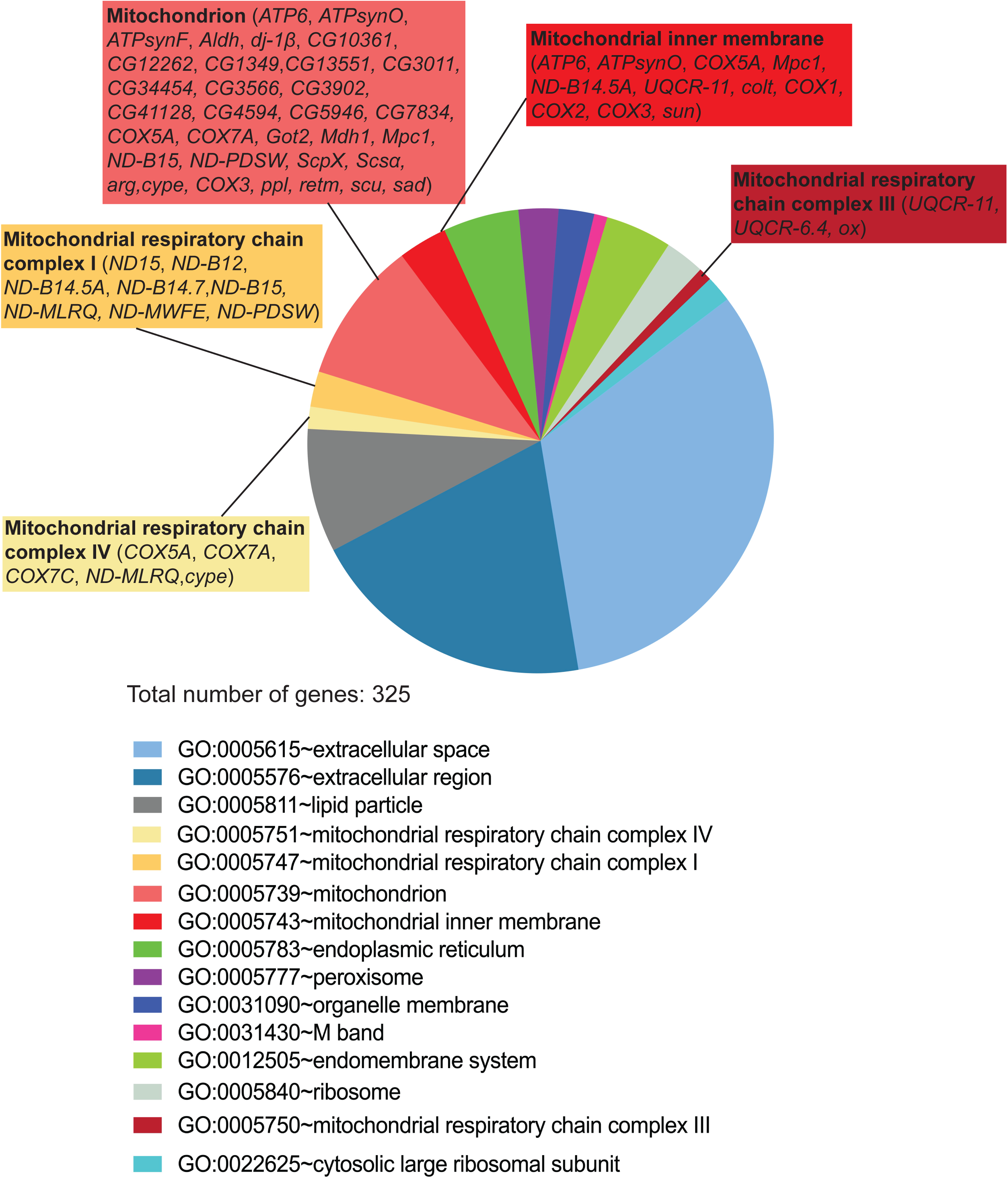
Gene ontology analysis for cellular component of genes specifically downregulated in aged, FHV-infected flies, reveal enrichment for mitochondria and mitochondrial respiratory chain complexes. GO analysis of at least two-fold differentially regulated genes 48h post FHV infection in aged flies. All identified categories are shown.

### Non-coding RNAs are differentially regulated by aging and FHV infection

We took a closer look at the differentially regulated genes, which were labeled as ‘other’ in our GO analysis (Figure S4). We observed that most of these genes are uncharacterized (categorized as candidate genes, or CG); several are non-coding RNAs (ncRNA); and others have previously described function but do not fit a specific DAVID GO category. Among ncRNAs, long non-coding RNAs (lncRNAs) correspond to a class of transcripts, which are at least 200nt long and lack a significant open reading frame (reviewed in (Sun and Kraus 2015)). Most lncRNAs are polyadenylated and can be reliably identified in our RNA-Seq workflow, in which an oligo-dT-based enrichment of poly-A-containing transcripts was used. A class of lncRNAs corresponds to antisense (as) RNAs, which are natural antisense transcripts (NATs) that overlap with protein-coding *loci* in the antisense direction.

We compared the number of ncRNAs differentially regulated at least two-fold in our RNA-Seq dataset and observed changes in expression of higher number of ncRNAs genes in aged FHV-infected than in young FHV-infected flies (68 vs 42 genes 24h p.i. and 267 vs 111 genes 48h p.i.) (Figure 6A, B and Figure S8A). Aging itself regulated the expression of 202 ncRNA genes. As observed for the total number of transcripts, ncRNAs, which were regulated by infection shared minimal overlap with aging (Figure 6B and Figure S8A). Among ncRNAs, we identified the largest proportion to correspond to lncRNAs. For all experimental groups we also found asRNAs and small nucleolar RNAs (snoRNAs). In young FHV-infected flies, a small percentage of ncRNAs corresponded to stable intronic sequence RNAs (sisRNAs). Specifically, in aged, FHV-infected flies we found differential regulation of ncRNAs that belong to small nuclear (snRNAs) and small non-messenger RNAs (snmRNAs) (Figure S8B).

**Figure 6.**
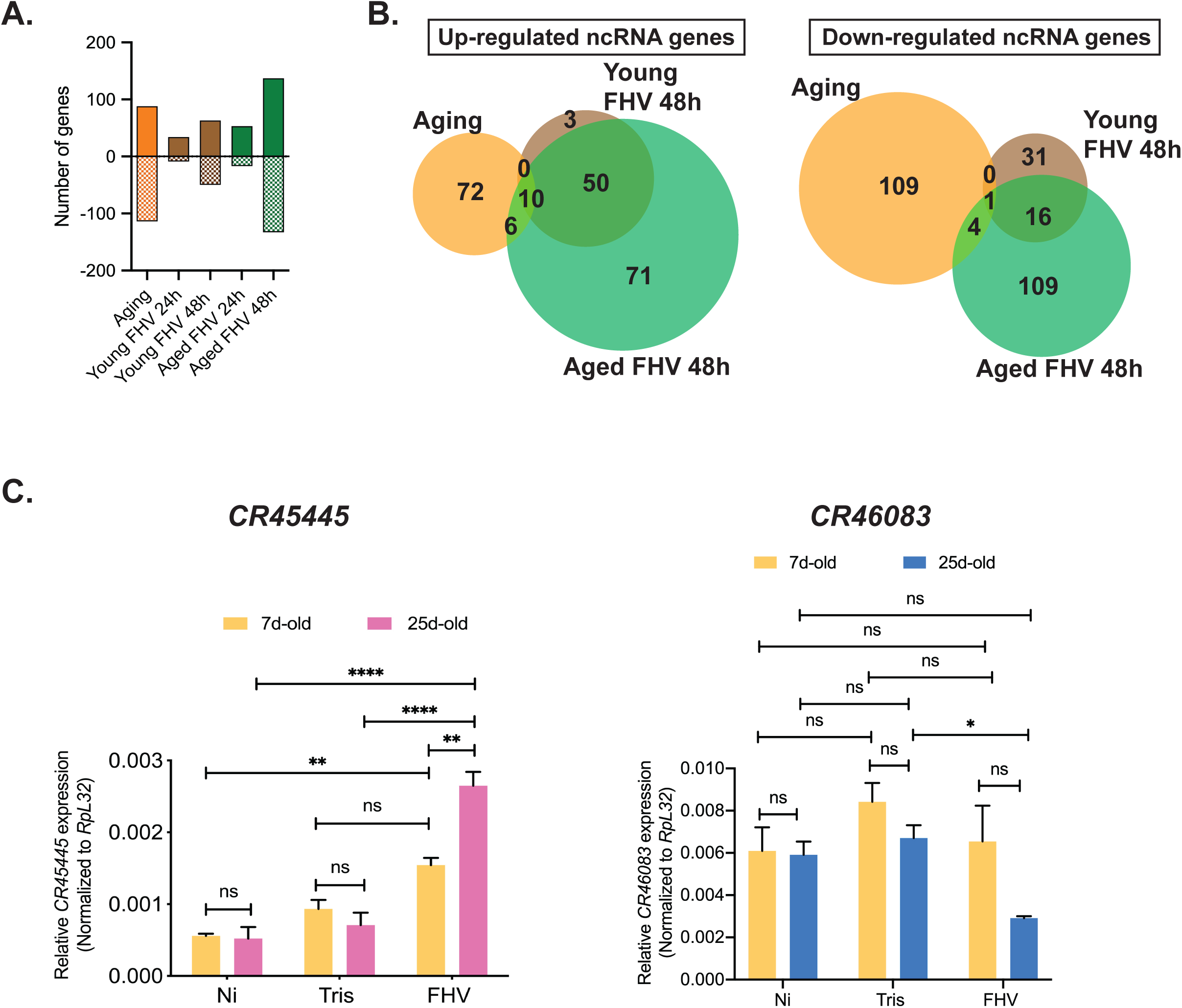
ncRNAs are differentially regulated by aging and infection. **A**. Comparison of the number of differentially regulated genes encoding for ncRNAs at least two-fold in all conditions. Positive values represent upregulated genes and negative values represent downregulated genes. **B**. Venn diagrams showing overlaps between differentially regulated ncRNA genes for selected experimental conditions. **C**. RT-qPCR-based gene expression analysis of asRNA *CR45445* and lncRNA *CR46083* 48h p.i. Statistics are based on two-way ANOVA followed by Tukey post-test to correct for multiple comparisons. ****: *p*<0.0001, ** : *p*<0.001, *: *p*<0.05, ns=non-significant.

We compared the expression of *CR45445* (an asRNA) and *CR46083* (an lncRNA) genes 48h p.i. by RT-qPCR. Consistent with the RNA-Seq data, we observed significant increase in *CR45445* and significant decrease in *CR46038* expression in comparison to Tris-injected controls in aged, but not young flies (Figure 6C). Together, these results indicate that both, aging and FHV infection affect the expression of genes encoding different categories of ncRNAs, and that specific ncRNAs are regulated in the aged organism after FHV infection.

## Discussion

We used the highly tractable genetic model *Drosophila melanogaster* to investigate the response of the aged organism following infection with the RNA(+) virus FHV. We found that 30-day old flies died faster than younger flies to FHV infection and that older, but not younger males were more sensitive than females. Although for both sexes we did not observe a difference in virus load as a function of age, our results indicate higher FHV titers in younger males in comparison to younger females, for which survival curves overlap. Although we cannot exclude genetic background-specific effects, our results raise the interesting question of whether control of virus replication in the young organism represents a sexually dimorphic trait. We observed that older males die faster than older females and contain twice the level of *FHV RNA1* transcript than females. This could potentially indicate that in comparison to females, younger males are able to tolerate higher FHV loads, but that this ability becomes impaired with age and results in more rapid death. Indeed, it is increasingly recognized that sexual dimorphism in immune function exists in *Drosophila* although the precise mechanisms underlying these age-dependent dimorphic differences are poorly understood (reviewed in (Belmonte *et al*. 2019)). More work is needed to elucidate this important aspect of immunity.

Both resistance and tolerance are components of host immunity (Martins *et al*. 2019). Antiviral RNAi is the main resistance mechanism that defends *Drosophila* against a broad range of RNA and DNA viruses, including FHV (Kemp *et al*. 2013). RNAi pathway mutants such as *Dicer-2* mutants, are more sensitive to FHV infection and mortality in *Dicer-2* mutants is accompanied by higher virus loads (Galiana-Arnoux *et al*. 2006). In this study, we find comparable FHV titers between young and aged flies in both whole bodies and circulating hemolymph. This suggests that aging likely affects a tolerance mechanism instead of resistance mechanisms. Earlier studies demonstrated that older *Drosophila* exhibit higher mortality following infection with *E*.*coli*, but were able to clear bacteria at similar rates as young flies (Ramsden *et al*. 2008) despite age-associated decline in macrophage function (Mackenzie *et al*. 2011; Horn *et al*. 2014). Both humoral (e.g. induction and secretion of AMPs) and cellular (e.g. phagocytosis) responses are required for bacterial clearance. The age-dependent increase in AMP expression could possibly compensate for decreased phagocyte function and account for the absence of an increase in bacterial load. Thus, the increased mortality following bacterial infection likely relies on age-dependent defects in tolerance. In our transcriptomic analysis we do not find noticeable transcriptional changes in gene expression of RNAi pathway components with aging, at least when flies are aged up to 25 days. This indirectly supports the hypothesis that antiviral RNAi is not functionally impaired in the aged fly. However, additional studies including small RNA sequencing during aging to compare the abundance of siRNAs against the FHV genome, are needed to determine whether this is the case.

We cannot entirely rule out the possibility that aging impacts resistance mechanisms in a tissue-specific way, differences which cannot necessarily be detected by measuring virus load in whole flies. It therefore would be very informative to perform additional studies to determine whether FHV differentially targets tissues at different ages and whether FHV load differs among tissues as a function of age. For instance, it is appreciated that aging affects gene expression differently in different tissues and in mammalian models differentially expressed genes in a given tissue are often not genes specific to this tissue (Rodwell *et al*. 2004). In *Drosophila*, a temporal and spatial transcriptional study of aging done on seven different tissues identified that <10% of differentially expressed genes in each tissue were in common with any other tissue (Zhan *et al*. 2007). It is therefore possible that host factors required for virus tissue tropism at younger age (e.g. in the heart and fat body (Eleftherianos *et al*. 2011)) become expressed in a different tissue in the aged host leading to shift in virus tropism accompanied by increased mortality even in the absence of higher virus titers. The aged *Drosophila*-FHV system could therefore represent an excellent model to address these questions and further examine how the aged organism is affected in the course of virus infection.

One striking finding of this study is that aged flies infected with FHV mount a more robust transcriptional response than younger flies. The fact that at 48h after FHV infection we find an overlap between 93% of upregulated genes and 57% of downregulated genes in young flies with genes regulated in aged flies, suggests that most of the transcriptional response to FHV is maintained as a function of age. However, aged flies show extensive regulation of additional genes. One possibility was that these additional genes are related to the process of aging itself. We show, however, that the overlap between the transcriptional profiles of aging, non-infected flies and aged, FHV-infected flies is minimal. The observed difference can potentially account for the changes in tolerance with age. Approximately three times more genes are downregulated than upregulated in aging flies in absence of viral infection. In aged, FHV-infected flies we see the opposite: a higher number of upregulated than downregulated genes for both time points examined. Thus, compared to younger adults, the aged fly mounts somehow a distinct response following FHV infection that is the consequence of the response to the virus rather than the process of aging itself. It remains unclear what factors contribute to the stronger transcriptional signature seen in aged FHV-infected flies. One hypothesis is that this possibly results from regulation by ncRNAs, including lncRNAs, which can play a role in transcriptional activation (Sun and Kraus 2015). Indeed, we find several lncRNAs regulated by infection specifically in aged, FHV-infected flies. Future experiments could address the question of whether lncRNAs, regulate the larger transcriptional response to FHV seen in aged flies.

Our study finds that the gene encoding the NF-kB transcription factor Relish as well as additional core components of the IMD pathway such as the adaptor protein Imd and the IKK complex component Key (IKKγ) are upregulated following FHV infection in aged flies. Interestingly, we find several AMP and IM genes that normally are upregulated in NF-kB - dependent way upon bacterial and fungal infections, to be downregulated following FHV infection even in aged flies. The role of NF-kB pathways in *Drosophila* antiviral immunity is complex and still not fully elucidated; however the pattern of expression that we see here aligns with previous findings, where infection with the DNA virus IIV-6 leads to downregulation of AMP genes, despite intact cleavage and nuclear translocation of Relish (West *et al*. 2019). Repression of AMP gene expression following IIV-6 infection appears downstream of Relish and likely occurs at the level of Relish binding to the AMP gene promoter or at the level of transcriptional activation (West *et al*. 2019). Whether in the case of FHV infection in aged flies the strong AMP and IM gene repression is mediated by similar mechanisms will remain a focus of future research.

Our results indicate that aged flies strongly upregulate *dSTING* expression in response to FHV 48h p.i. In young flies, dSTING does not play a protective role against FHV, as *dSTING* null mutants show similar, if not slightly better, survival to FHV in comparison to controls (Goto *et al*. 2018). It may be that in response to FHV, dSTING in aged flies plays a pro-death, rather than pro-survival role. In addition to its essential role in interferon production, STING signaling in mammals plays a role in the activation of programmed cell death, including Caspase-9 and Caspase-3-mediated apoptosis, although the exact mechanisms are not well understood (reviewed in (Maelfait *et al*. 2020)). Thus, it is possible that in response to FHV, dSTING mediates the strong apoptotic signature, that could be associated with the more rapid death observed in aged flies. Future analysis of *dSTING* function in older flies in response to FHV could for instance reveal novel information about evolutionary conservation of dSTING-mediated apoptotic signaling. Additionally, because of increased apoptotic gene deregulation and the fact that phagocytic function decreases with age, future experiments should be also aimed at examining whether defective apoptotic corpse clearance is associated with the higher mortality of older flies following FHV infection.

Our transcriptomic analyses reveal that as FHV infection progresses in aged flies, genes associated with mitochondrial respiratory chain become downregulated. Additionally, we notice that several transcripts of genes encoded by the mitochondrial genome (Table S3) are detected in the Aged FHV samples, especially at the early, 24h time point, suggesting a link to apoptosis. One possible scenario is that p53-mediated apoptotic cell death is activated early in response to FHV leading mitochondria to become leaky and to release transcripts of genes that are encoded by the mitochondrial genome. Induction of pro-apoptotic gene expression within the first hours immediately following FHV infection of adult flies is an important mechanism that limits virus replication (Liu *et al*. 2013). It is important to examine the dynamics of this response in young and aged flies to determine whether any differences in this very early response are present between the two age groups. FHV-triggered apoptosis in the aged fly can also account for the downregulation of genes involved in the ETC and generation of ATP. As a consequence, it is possible that the bioenergetic profile of the cell is reduced and mitochondrial respiration halted post-infection in aged flies. Because programmed cell death and ATP production are increasingly considered closely linked aspects of mitochondrial function (Burke 2017), it will be important for future studies to determine whether FHV triggers apoptosis-dependent changes in cellular bioenergetics and how this relates to the more rapid death of the aged, FHV-infected organism.

In conclusion, in this study we addressed for the first time how aged *Drosophila* respond to infection with the plus-strand RNA virus FHV and provide a detailed transcriptional comparison of the responses between young and aged flies at two time points following infection. With the advantages that *Drosophila* offer to investigate gene function, this study sets up the stage for future investigations about the mechanisms that underlie aged host-virus interactions using not only FHV, but also other viruses. For instance, DCV triggers distinct pathophysiological events in comparison with FHV (Chtarbanova *et al*. 2014), and it also leads to the more rapid death of older flies (Eleftherianos *et al*. 2011). It would be very interesting to explore the age-dependent response to DCV infection, as this could lead to the discovery of additional mechanisms that help the aged organism survive virus infection.

### Experimental procedures

#### Drosophila handling

All *Drosophila* stocks were raised and maintained on Nutri-Fly_®_ Bloomington formulation food (Genesee Scientific, Cat #: 66-113) at 25°C. *Oregon-R* (#2376) and *y*_*1*_ *w* _*67c23*_ (#6599) flies were obtained from the Bloomington *Drosophila* Stock Center (Bloomington, IN). *w*_*1118*_ flies were a kind gift from Dr. John Yoder (University of Alabama). For aging experiments, 0-4 days-old animals were collected, CO_2_-anesthetized, separated by sex and placed in a 25°C incubator with controlled 12/12 dark/light cycle. Flies were flipped every two to three days in a fresh food-containing vial until desired age was reached. For survival and virus load determination young flies were 3-7-day old (labeled as 5d-old), and aged flies were 27-31-day old (labeled as 30d-old). For RNA-Seq experiments, replicates containing young flies were 6-9 days-old (labeled as 7d-old), and aged flies were 22-29 days-old (labeled as 25d-old), Table S6. *Wolbachia*-free flies were used in all experiments.

### Virus stock and infections

Flock House Virus (FHV) was a kind gift from Dr. Annette Schneemann (Scripps Research Institute, La Jolla, CA). FHV stock titer was determined at 2.92E+06 TCID50/mL (Figure S9) using the method as in (Eleftherianos *et al*. 2011). Flies of desired sex, age and genotype were individually injected with 4.6nL of either virus stock solution or control 10mM Tris-HCl pH7.5 solution under CO_2_ anesthesia using a Nanoject II injector (Drummond Scientific). Flies were let to recover from the injection for ∼one hour at room temperature and then were placed in a 22°C incubator. For survival experiments, flies were separated by sex and placed in groups of 10 per vial for each experimental treatment. The number of living flies was recorded every 24 h. For virus load determination by RT-qPCR, flies were separated by sex and frozen in groups of 5 flies per experimental treatment prior RNA extraction.

### RNA sequencing

The Quick-RNA MiniPrep Kit (Zymo Research) was used to isolate total RNA from fifteen whole flies. Three biological replicates were collected for each experimental condition. RNA was extracted following manufacturer’s instructions and sent to Novogene Co., Ltd. for RNA sequencing. Prior directional library preparation, quality of RNA for all samples was evaluated by Novogene Co., Ltd. for purity, degradation, potential contamination and integrity. Only for samples that passed quality control mRNA was enriched using oligo(dT) beads. Constructed libraries were quality checked and paired end sequencing performed using Illumina technology. Initial bioinformatics analysis to determine differential gene expression was performed by Novogene Co., Ltd using the *Drosophila melanogaster* reference genome (dmel_r6.23_FB2018_04). Readcounts were normalized using the DESeq 1.10.1 (Anders and Huber 2010) method and adjusted *p*-values (*p adj*) estimated based on a negative binomial distribution model. *p adj* <0.05 were considered significant. Validation of gene expression by RT-qPCR was performed on RNA used for the RNAseq experiment. Determination of differential gene expression in experimental groups is as follows: Aging: Non-infected 25d/Non-infected 7d; Young FHV24h: 7d FHV 24h/7d Tris 24h; Aged FHV24h: 25d FHV 24h/25d Tris 24h. Young FHV48h: 7d FHV 48h/7d Tris 48h; Aged FHV48h: 25d FHV 48h/25d Tris 48h.

### RT-qPCR gene expression analysis

The Quick-RNA MiniPrep Kit (Zymo Research) was used to isolate total RNA following manufacturer’s instructions. RNA (1000ng) was converted to cDNA using the High Capacity RNA-to-cDNA Kit (Applied Biosystems). RT-qPCR reaction was performed using *Power* SYBR™ Green PCR Master Mix (Applied Biosystems) according to manufacturer’s instructions. Primer sequences are listed in Table S7. For all assays, expression of *RpL32* (*Rp49)* was used to normalize gene expression.

### Statistical analysis

Statistical analysis of median survival, virus load and gene expression analysis were performed using GraphPad Prism 8 for MAC software and *p* < 0.05 considered significant.

### Functional annotation analysis

We used the Database for Annotation, Visualization and Integrated Discovery (DAVID) 6.8 (Huang *et al*. 2009b; Huang *et al*. 2009a) to analyze enriched functional gene categories, including gene ontology (GO) and KEGG pathways for differentially regulated genes at least two-fold. The cut-off *p* value to determine enriched GO categories and pathways was set at 0.1.

## Data availability

Raw sequencing reads generated during this project have been deposited with the National Center for Biotechnology Information Sequence Read Archive under BioProject PRJNA644593. File names corresponding to experimental samples are shown in Table S7. The authors affirm that all data necessary for confirming the conclusions of the article are present within the article, figures, and tables, and in supplemental material. Supplemental files including supplemental experimental procedures, figures and tables have been deposited to the GSA Figshare portal. Supplemental tables S3 and S5 have been submitted as Excel files while all other supplemental materials are in a PDF format.

## Acknowledgements

We thank Dr. Annette Schneemann for FHV virus stock and the anti-FHV antibody used in this study. We are grateful to Drs. David Wassarman and Grace Boekhoff-Falk for critical reading of the manuscript. The Authors declare that they have no conflict of interest.

## Author contributions

SC conceptualized the study and designed experiments; LS, NMS, AE, EH, MD and CG performed experiments and data collection; SC, LS, NMS, AE, EH, CG and MD analyzed data, JLF analyzed RNA-Seq data; SC and JLF wrote the manuscript with input from all authors. All authors red and approved the manuscript.

## References

Anders, S., and W. Huber, 2010 Differential expression analysis for sequence count data. Genome Biol 11: R106.

Belmonte, R. L., M. K. Corbally, D. F. Duneau and J. C. Regan, 2019 Sexual Dimorphisms in Innate Immunity and Responses to Infection in Drosophila melanogaster. Front Immunol 10: 3075.

Boyle, M., C. Wong, M. Rocha and D. L. Jones, 2007 Decline in self-renewal factors contributes to aging of the stem cell niche in the Drosophila testis. Cell Stem Cell 1: 470–478.

Burke, P. J., 2017 Mitochondria, Bioenergetics and Apoptosis in Cancer. Trends Cancer 3: 857–870.

Cao, Y., S. Chtarbanova, A. J. Petersen and B. Ganetzky, 2013 Dnr1 mutations cause neurodegeneration in Drosophila by activating the innate immune response in the brain. Proc Natl Acad Sci U S A 110: E1752–1760.

Carney, T. D., A. J. Struck and C. Q. Doe, 2013 midlife crisis encodes a conserved zinc-finger protein required to maintain neuronal differentiation in Drosophila. Development 140: 4155–4164.

Chtarbanova, S., O. Lamiable, K. Z. Lee, D. Galiana, L. Troxler et al., 2014 Drosophila C virus systemic infection leads to intestinal obstruction. J Virol 88: 14057–14069.

Eleftherianos, I., S. Won, S. Chtarbanova, B. Squiban, K. Ocorr et al., 2011 ATP-sensitive potassium channel (K(ATP))-dependent regulation of cardiotropic viral infections. Proc Natl Acad Sci U S A 108: 12024–12029.

Fabian, D. K., K. Garschall, P. Klepsatel, G. Santos-Matos, E. Sucena et al., 2018 Evolution of longevity improves immunity in Drosophila. Evol Lett 2: 567–579.

Franceschi, C., M. Bonafe, S. Valensin, F. Olivieri, M. De Luca et al., 2000 Inflamm-aging. An evolutionary perspective on immunosenescence. Ann N Y Acad Sci 908: 244–254.

Galiana-Arnoux, D., C. Dostert, A. Schneemann, J. A. Hoffmann and J. L. Imler, 2006 Essential function in vivo for Dicer-2 in host defense against RNA viruses in drosophila. Nat Immunol 7: 590–597.

Garschall, K., and T. Flatt, 2018 The interplay between immunity and aging in Drosophila. F1000Res 7: 160.

Goto, A., K. Okado, N. Martins, H. Cai, V. Barbier et al., 2018 The Kinase IKKbeta Regulates a STING-and NF-kappaB-Dependent Antiviral Response Pathway in Drosophila. Immunity 49: 225–234 e224.

He, W., D. Goodkind and P. Kowal, 2016 An Aging World: 2015., pp. in U.S. Census Bureau, International Population Reports.

Horn, L., J. Leips and M. Starz-Gaiano, 2014 Phagocytic ability declines with age in adult Drosophila hemocytes. Aging Cell 13: 719–728.

Huang, D. W., B. T. Sherman and R. A. Lempicki, 2009a Bioinformatics enrichment tools: paths toward the comprehensive functional analysis of large gene lists. Nucleic Acids Research 37: 1–13.

Huang, D. W., B. T. Sherman and R. A. Lempicki, 2009b Systematic and integrative analysis of large gene lists using DAVID bioinformatics resources. Nature Protocols 4: 44–57.

Kemp, C., S. Mueller, A. Goto, V. Barbier, S. Paro et al., 2013 Broad RNA interference-mediated antiviral immunity and virus-specific inducible responses in Drosophila. J Immunol 190: 650–658.

Kounatidis, I., S. Chtarbanova, Y. Cao, M. Hayne, D. Jayanth et al., 2017 NF-kappaB Immunity in the Brain Determines Fly Lifespan in Healthy Aging and Age-Related Neurodegeneration. Cell Rep 19: 836–848.

Lamiable, O., and J. L. Imler, 2014 Induced antiviral innate immunity in Drosophila. Curr Opin Microbiol 20: 62–68.

Lemaitre, B., and J. Hoffmann, 2007 The host defense of Drosophila melanogaster. Annu Rev Immunol 25: 697–743.

Leng, J., and D. R. Goldstein, 2010 Impact of aging on viral infections. Microbes Infect 12 : 1120–1124.

Liu, B., S. K. Behura, R. J. Clem, A. Schneemann, J. Becnel et al., 2013 P53-mediated rapid induction of apoptosis conveys resistance to viral infection in Drosophila melanogaster. PLoS Pathog 9: e1003137.

Liu, Y., B. Gordesky-Gold, M. Leney-Greene, N. L. Weinbren, M. Tudor et al., 2018 Inflammation-Induced, STING-Dependent Autophagy Restricts Zika Virus Infection in the Drosophila Brain. Cell Host Microbe 24: 57–68 e53.

Mackenzie, D. K., L. F. Bussiere and M. C. Tinsley, 2011 Senescence of the cellular immune response in Drosophila melanogaster. Exp Gerontol 46: 853–859.

Maelfait, J., L. Liverpool and J. Rehwinkel, 2020 Nucleic Acid Sensors and Programmed Cell Death. J Mol Biol 432: 552–568.

Martins, R., A. R. Carlos, F. Braza, J. A. Thompson, P. Bastos-Amador et al., 2019 Disease Tolerance as an Inherent Component of Immunity. Annual Review of Immunology, Vol 37, 2019 37: 405-437.

Merkling, S. H., G. J. Overheul, J. T. van Mierlo, D. Arends, C. Gilissen et al., 2015 The heat shock response restricts virus infection in Drosophila. Sci Rep 5: 12758.

Mussabekova, A., L. Daeffler and J. L. Imler, 2017 Innate and intrinsic antiviral immunity in Drosophila. Cell Mol Life Sci 74: 2039–2054.

Nikolich-Zugich, J., 2018 The twilight of immunity: emerging concepts in aging of the immune system. Nat Immunol 19: 10–19.

Nikolich-Zugich, J., K. S. Knox, C. T. Rios, B. Natt, D. Bhattacharya et al., 2020 SARS-CoV-2 and COVID-19 in older adults: what we may expect regarding pathogenesis, immune responses, and outcomes (vol 42, pg 505, 2020). Geroscience 42: 1013–1013.

Pletcher, S. D., S. J. Macdonald, R. Marguerie, U. Certa, S. C. Stearns et al., 2002 Genome-wide transcript profiles in aging and calorically restricted Drosophila melanogaster. Curr Biol 12: 712–723.

Ramsden, S., Y. Y. Cheung and L. Seroude, 2008 Functional analysis of the Drosophila immune response during aging. Aging Cell 7: 225–236.

Rodwell, G. E., R. Sonu, J. M. Zahn, J. Lund, J. Wilhelmy et al., 2004 A transcriptional profile of aging in the human kidney. PLoS Biol 2: e427.

Settles, E. W., and P. D. Friesen, 2008 Flock house virus induces apoptosis by depletion of Drosophila inhibitor-of-apoptosis protein DIAP1. J Virol 82: 1378–1388.

Shah, P. S., N. Link, G. M. Jang, P. P. Sharp, T. Zhu et al., 2018 Comparative Flavivirus-Host Protein Interaction Mapping Reveals Mechanisms of Dengue and Zika Virus Pathogenesis. Cell 175: 1931–1945 e1918.

Sun, M., and W. L. Kraus, 2015 From discovery to function: the expanding roles of long noncoding RNAs in physiology and disease. Endocr Rev 36: 25–64.

Tatar, M., 2010 Reproductive aging in invertebrate genetic models. Ann N Y Acad Sci 1204: 149–155.

Venter, P. A., and A. Schneemann, 2008 Recent insights into the biology and biomedical applications of Flock House virus. Cell Mol Life Sci 65: 2675–2687.

West, C., F. Rus, Y. Chen, A. Kleino, M. Gangloff et al., 2019 IIV-6 Inhibits NF-kappaB Responses in Drosophila. Viruses 11.

Zerofsky, M., E. Harel, N. Silverman and M. Tatar, 2005 Aging of the innate immune response in Drosophila melanogaster. Aging Cell 4: 103–108.

Zhan, M., H. Yamaza, Y. Sun, J. Sinclair, H. Li et al., 2007 Temporal and spatial transcriptional profiles of aging in Drosophila melanogaster. Genome Res 17: 1236–1243.

